# Nanomechanical and microrheological properties of bladder cancer cells at cellular and spheroid levels

**DOI:** 10.1101/2022.03.21.485153

**Authors:** Kajangi Gnanachandran, Sylwia Kędracka-Krok, Joanna Pabijan, Małgorzata Lekka

## Abstract

Cancer progression is associated with changes in cell mechanical and rheological properties that could be probed by atomic force microscopy (AFM). In this study, we applied AFM to measure elastic (by compressing the cells) and viscoelastic (by applying shear stress) properties of bladder cancers in relation to their culture morphology (in single cells, cell monolayers, and spheroids). Three different cell lines, HCV29 (non-malignant cell cancer of ureter), HT1376 (grade III bladder carcinoma), T24 (grade IV transitional cell carcinoma), were investigated. Nanoindentation measurements only differentiate between non-malignant and cancer cells, but it is difficult to distinguish between specific bladder cancers. By applying microrheological measurements, we confirm that non-malignant cells are more rigid than cancer cells but more importantly, it was possible to differentiate between two cancerous cell lines, regardless of the culture conditions. As each of them is characterized by a distinct actin filament network inside the cell, we showed that actin filaments are a key element in defining the rheological properties of spheroids originating from cells having thick actin bundles. Our results showed that HCV29 cells are more rigid than the studied cancer cells, indicating that normal cells are resistant to compressive and shear forces. Therefore, we conclude that cell mechanical and rheological properties may serve as a biophysical marker to distinguish normal and cancer cells of different malignancies.

## Introduction

Bladder cancers are among the most common malignancies known to cause cancer death worldwide. Almost 90% of this type of cancer is caused by the urothelial cells (transitional epithelial cells) lining the bladder and urinary tract. These cells are highly exposed to mutagenic agents sent from the kidney into the urine^1,2^. Cancer progression is associated with genetic mutations that change key functions of the cells, such as cellular growth, differentiation, interactions with the extracellular matrix (ECM) and the adjacent cell^3^. All these features enable replicative immortality and the activation of invasion and metastasis^4^. Oncogenic transformation is related to changes in the mechanical/rheological properties of cells^5,6^ and followed by rearrangements in actin cytoskeleton^7,8^. In tumours, tensional tissue homeostasis is altered due to differences in rheological properties of cells and increased cell-generated forces^9^. This also comes from the fact that the cells *in vivo* grow next to each other. Thus, they receive and send signals from and to the neighbouring cells. Cell-cell communication can be mediated in different ways by the interaction of signaling molecules such as growth factors, hormones, chemokines, and cytokines: (1) paracrine cell-cell communication, which does not require the physical contact between cells, (2) juxtacrine communication where cells directly exchange the signaling molecules without secreting them into the extracellular space and (3) Endocrine communication is typically mediated by hormones that travel long distances through extracellular fluids^10^. During cancer progression, cell signaling is deregulated to resist pro-apoptotic pathways and invade the surrounding tissues^11^. Therefore, the single non-confluent cells or the cells forming a monolayer or spheroids may exhibit different properties when cultured in vitro, and these may cause the rearrangement of the actin cytoskeleton^12^. Nowadays, 3D spheroids have been widely applied as tumour models because they are used to model and mimic the natural environment of cells and some important features of solid tumours, such as cell-cell signaling, cell-matrix interactions, and drug resistance mechanism^13^. However, there are only a few available pieces of information regarding their physical characterization, especially for bladder cancer. Therefore, in this study, we compare the mechanical and microrheological properties of bladder cancer cells in 2D cultures and in 3D spheroids.

Atomic force microscopy (AFM) is a versatile and powerful tool that unravels the relation between cellular structures and mechanical properties^14–16^. The technique has been used to study the elastic properties of cells originating from various cancers such as ovarian, breast, prostate, and thyroid^17–21^. Several AFM experiments have been performed on bladder cancer cells^22–26^. Cancerous cells have consistently been softer than normal or benign cells in all these studies, showing a lower elastic (Young’s) modulus value. These changes have been linked with reorganizing the cell cytoskeleton, especially the actin filaments^7,22,27^. Cells are a complex viscoelastic material^28^ composed of both elastic and viscous contributions^29^. The viscoelastic nature of cancerous cells has been demonstrated to be a valuable marker of their metastatic potential^30–32^. For this reason, it is advisable to measure not only their elastic but also viscous properties in parallel. Few studies were carried out by considering the effects of load rate in the context of the indentation experiments bringing the direct experimental relation between the measured mechanical properties and the applied load rate^18,33,34,35^. Other AFM-based methods measuring cell viscoelasticity apply the sinusoidal oscillations of the cantilever with a small and constant amplitude using a broad frequency range^36,37^. Such experiments require theoretical models describing the viscoelastic nature of the cells. The power-law rheological model is among the most widely applied ones. This model stems from the soft glassy rheology theory^38^, and it suggests that the contraction and remodelling of the cytoskeleton is the reason behind the rheological behaviour of the cells^36,39^. So far, there have been only a few research works in which AFM characterized the physical properties of the spheroids, and they were all focused on studying the mechanical properties and not the rheology in details^40–44^.

Here, we focused on the mechanical (compressive) and rheological properties of bladder cancer cells at three levels of cellular organization. We compared cell properties for single cells, cell monolayers (referred to as 2D cultures), and spheroids composed of either from HCV29 (non-malignant cancerous cells) or T24 (grade IV transitional cell carcinoma cells) or HT1376 (grade III carcinoma cell) cells. Our previous studies^25,27,45^ show that it is easy to measure the difference between non-malignant and cancerous cells. However, distinguishing between various bladder cancer types is not easy, especially from data collected during indentation. That is why we investigate whether rheological parameters are better for differentiating the non-malignant cells from cancer cells and allowing to distinguish different bladder cancer types. In parallel, we focused on actin filaments as major cytoskeletal elements responsible for cell resistance to deformations. We hypothesize that their contribution is a major component of the cellular mechanical/rheological response regardless of the cell organization level.

## Materials and methods

### Cell cultures

Three human bladder cancer cell lines were used in this study: HCV29 – non-malignant transitional epithelial cancer cell of the ureter (obtained from the Institute of Experimental Therapy, PAN, Wroclaw, Poland), T24 – transitional cell carcinoma (ATCC, LGC Standards) and HT1376 – grade III urinary bladder cell carcinoma (ATCC, LGC Standards). HCV29 and T24 were cultured in RPMI-1640 medium (Sigma) supplemented with 10% of fetal bovine serum (FBS, Sigma). HT1376 were grown in Eagle’s minimum essential medium (EMEM, LGC Standards) with 10% FBS. All cell lines were grown in culture flasks (Sarstedt) in an incubator (Nuaire) at 37° C in 95% air and 5% CO2 atmosphere. The relative humidity was kept above 98%. The cell passaging was carried out when they reached 80-90% of the confluency level. For HCV29 and T24 cells, 0.05% and for HT1376 cells, 0.25% of trypsin-EDTA solution (Sigma) was applied. After a few passages (< 10), cells were ready to be seeded on a Petri dish or glass coverslips for fluorescence imaging and AFM measurements. *STR profile of HCV29 cell line*: amelogenin: X,Y; TH01: 6,8; D13S317: 8,11; D16S539: 12,13; D5S818: 12; D7S820: 7,12; TPOX: 8,12; vWA: 19,21; CSF1PO: 11,12; *STR profile of HT1376 cell line*: amelogenin: X; TH01: 7,10; D13S317: 9,11; D16S539: 11,14; D5S818: 11,12; D7S820: 9,12; TPOX: 8; vWA: 15,18; CSF1PO: 12; *STR profile of T24 cell line:* X; TH01: 6; D13S317: 12; D16S539: 9; D5S818: 10,12; D7S820: 10,11; TPOX: 8,11; vWA: 17,19; CSF1PO: 10,12;

### Preparation of cancer spheroids

Spheroids were prepared in 96-well U-bottom 3D cell culture plates (Thermofisher). The initial seeding density of cells/well was adjusted differently to the three cell lines set to obtain spheroids of 350-400 µm diameter after 3 days of incubation. They are 6000 (HCV29), 10000 (T24), and 1200 (HT1376) cells per well. For the AFM measurements, the spheroids were collected from the culture wells and placed in micromesh arrays with square holes made of bio-compatible silicone polymer, which were attached to the Petri dish. This helped to keep the spheroids immobile during the indentation.

### Determining G/F actin ratio

The G-actin/F-actin In Vivo Assay Kit (cat. #BK037, Cytoskeleton) was used to determine the F/G-actin ratio. It was determined in cells cultures as a monolayer. After homogenizing cells in 300 µl of LAS2 buffer (LAS2 contains detergents disrupting the cell membrane), which solubilizes G-actin and not F-actin, the cell lysate was centrifuged at 100.000 x g at 37°C for 1h, separating a soluble G-actin from the polymerized F-actin (ultracentrifuge was accessible at the Department of Physical Biochemistry, Faculty of Biochemistry, Biophysics and Biotechnology, Jagiellonian University). F-actin was present at the bottom of the pellet, while the supernatant contained G-actin.

### Western blot analysis

The Western blot was applied to analyse the G- and F-actin contents using the following protocol. The collected supernatants were separated on 10% SDS-PAGE gels and transferred onto a PVDF membrane. Anti-actin rabbit polyclonal antibody, provided along with the kit (Cytoskeleton) was used to detect G- and F-actins. Bands were visualized using horseradish peroxidase-coupled secondary anti-rabbit antibody (Cell Signaling Technology). The ratio between G/F actin was determined by using Image J based densitometric calculations.

### Cytochalasin-D treatment

Cells were grown in Petri dishes for the 2D cultures (single cells and monolayers) and in 96-well U-bottom plates used to form spheroids. Afterwards, the culture medium was removed, and phosphate-buffered saline (PBS, Sigma) containing 5 µM cytochalasin D (Sigma) was added for 30 minutes at 37°C. Then, the samples were carefully washed, and a fresh culture medium was added before the AFM measurements.

### Epi-fluorescence microscopy

Cells were cultured in Petri dishes until the desired density of cells was obtained, i.e., single cells or cell monolayers. Cells were washed with PBS buffer and fixed by adding a 3.7% paraformaldehyde solution in PBS for 20 minutes. Next, they were washed with PBS three times, and the cell membrane was permeabilized using 0.2% Triton X-100 (Sigma) in PBS at 4° C for 5 minutes. Afterwards, the cells, rinsed with PBS, were labelled with fluorescent dyes in an environment with a minimum light level. Phalloidin conjugated with Alexa Fluor 488 (Invitrogen) was dissolved in PBS buffer (1:200) and used to stain the actin filaments for 30 minutes, while the cell nuclei were stained with Hoechst 33342 (Invitrogen) dissolved in PBS buffer (1:5) for 15 minutes. Afterwards, the samples were washed with PBS buffer. The fluorescent images were recorded using an Olympus IX83 inverted microscope (Olympus, Japan) equipped with a 100 W mercury lamp and a set of filters used to record the emission at 515-545 nm (Alexa Fluor 488) and 461 nm (Hoechst 33342). The images were acquired using the Orca Spark digital camera (Hamamatsu, Japan), providing 2.3 megapixels (1920 pixels per 1200 pixels) images that were recorded using CellSens Dimension software (Olympus).

### Confocal microscopy

Confocal images of actin filaments inside spheroids were visualized using a confocal microscope, accessible in the Laboratory of in vivo and in vitro imaging, Maj Institute of Pharmacology, Polish Academy of Sciences. The samples were stained using the following protocol. Spheroids from each cell line were collected in a 1.5 ml Eppendorf and fixed with 3.7% paraformaldehyde for 1 hour. Following this, the samples were washed with PBS for 2 minutes three times, treated with 1% cold Triton X-100 overnight at 4°C, and washed again with PBS. Then, 1% cold Bovine Serum Albumin solution (BSA, Sigma) was added for 3 hours at 4°C, and after removing BSA, the spheroids were washed with PBS. Next, the spheroids were incubated overnight at 4°C with phalloidin conjugated with Alexa Fluor 488 (1:20 in PBS). The next day, the dye for actin filaments was removed, and SiR-DNA (Spirochrome), diluted 1:1000 in PBS, for the cell nuclei was added. Finally, the samples were washed with PBS and moved to 18-well glass-bottom slides (Ibidi) with an anti-shading solution (Thermofisher) having the same refractive index (1.52) as the oil used for the objective of 63 X magnification. Images were recorded using a Leica TCS SP8 WLL confocal microscope, equipped with new-generation HyD detectors set at 730-800 nm (SiR-DNA) and 509-560 nm (Alexa Fluor 488). Fluorescent dyes were excited by diode lasers: 405 nm (Hoechst) and white light laser with emission wavelength set at 499 nm (AlexaFluor 488). Images were registered using an oil immersion 63x objective lens (HC PL APO CS2 NA 1.40).

### Atomic force microscopy (AFM)

The mechanical and microrheological properties of cellular samples, i.e., 2D cultures and spheroids, were measured using two AFMs. A commercial AFM (Xe120 model, Park Systems, Korea) working in a force spectroscopy mode and integrated with an inverted optical microscope (Olympus IX71) was mainly applied for nanoindentation measurements, while Nanowizard IV (Bruker – JPK Instruments), also coupled to an inverted optical microscope (Olympus IX71) was applied to measure rheological properties using the MicroRheology module (**Fig. 1**).

**Fig. 1.**
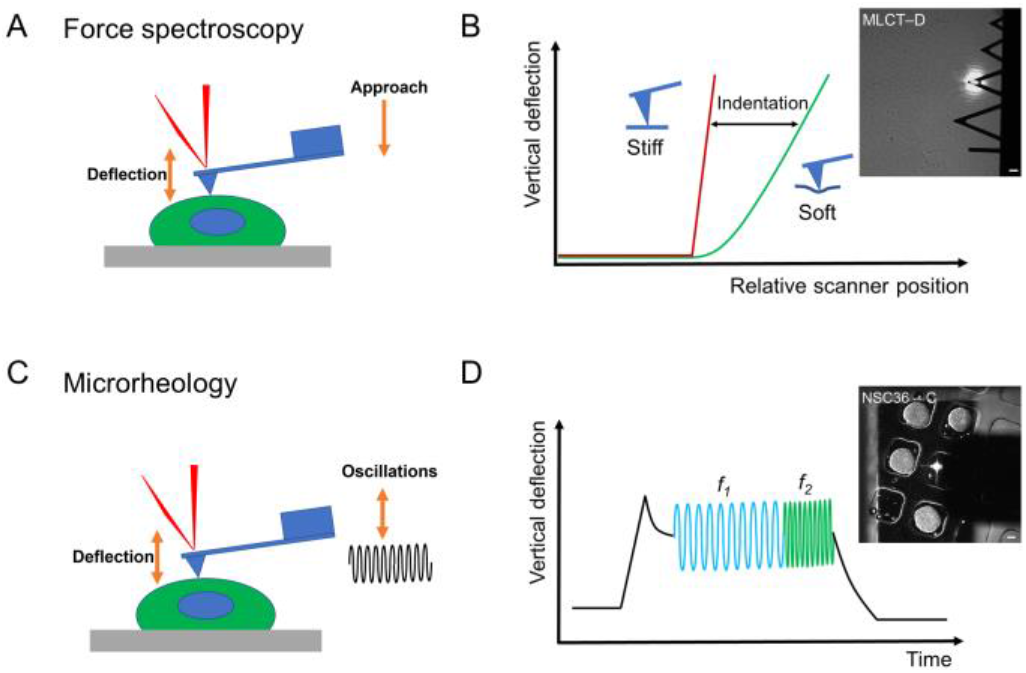
AFM-based nanomechanics and microrheology applied to assess mechanical and rheological properties of 2D cultures and spheroids. Nanoindentation (A) and microrheological (C) measurements. In nanoindentation (B), the difference in force curves recorded for rigid and compliant materials is used to quantify Young’s (elastic) modulus. In microrheological measurements (D), a sine modulation is applied at a certain indentation. The resulting oscillations in the vertical cantilever deflection served as a basis for shear modulus determination. During indentation, force curves were recorded over the nuclear region within the scan area of 64 µm^2^ (a map of 6 pixels per 6 pixels was set; 36 force curves were collected for single cells). In microrheological measurements, sine modulations were recorded for frequencies from 1 Hz to 100 Hz (2D cultures) and from 1 Hz to 250 Hz (spheroids), taken from the central cell or spheroid regions. Insets: Top view image of cells and spheroids measurements setup in the AFM. Two different cantilevers are shown MLCT-D (used for 2D cell cultures) and NSC36-C (used for spheroids).

Commercially available silicon nitride triangular cantilevers (MLCT-D, Bruker, USA) were used. They are characterized by a nominal spring constant of 0.03 N/m, frequency of 15 kHz, the width of 20 um, length of 225 um, tip radius of 20 nm. The tip was a four-sided pyramid described by the following angles: the front angle of 15 ± 2.5º, back angle of 25 ± 2.5º, and side angle of 17.5 ± 2.5º. An open angle of 20º was applied in the calculations (an average from all angles). The sensitivity of the photodetector was calibrated on a rigid glass surface. Spheroids were probed with sharp silicon nitride cantilevers (NSC36-C, Park Systems) characterized by a spring constant of 0.6 N/m (resonant frequency of 65 kHz). They possess a pyramidal probe with a full open-angle of 40º (a half open-angle equals 20º). Cantilever spring constants were calibrated using the thermal tune method^46,47^. The exact values were 0.036 ± 0.004 N/m (mean ± standard deviation, SD, from n = 8) and 0.59 ± 0.04 N/m (n = 3) for MLCT-D and NSC36-C cantilevers, respectively.

### AFM-based nanoindentation

Force curves were acquired within a region of 64 µm^2^ above the cell nuclear region (or in the central part of the spheroid), with a 6 pixel × 6 pixel map (in total, 36 curves were recorded per cell). The approach speed was set to 8 µm/s, and the load force was kept below 1-2 nN. For each set of experiments conducted for 2D cultures, 20 cells were measured in liquid conditions (i.e., cells in the corresponding culture medium) at RT, no longer than two hours. The measurements were repeated three times. In the case of spheroids measured at the analogous liquid conditions, 15 spheroids for each cell line were measured (2 force maps per single spheroid). A scan area of 9 µm^2^ was chosen (a grid of 3 pixels per 3 pixels was set in the central part of the spheroid). In total, 18 force curves were recorded per single spheroid. Force curves were collected with the approach speed of 8 µm/s and load force of 15 nN. Measurements were repeated three times.

### Young’s modulus determination

The apparent Young’s (elastic) modulus was determined by applying the Hertz-Sneddon contact mechanics^48^. The force curves were recorded on a stiff, non-deformable surface (calibration curve recorded here on a glass coverslip) and on soft samples (cells or spheroids). Next, the calibration curves were subtracted from those recorded on samples (**Fig. 1 D**).

The obtained force versus indentation curve was fitted with the equation relating the load force *F* and indentation *δ*, assuming a conical shape of the probing pyramidal tip:

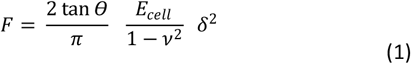

where *Θ* is the half open-angle of the cone, *ν* is the Poisson ratio of the cell (set to 0.5 treating cells as incompressible materials), and *Ecell* is the apparent Young’s modulus. Calculations were performed for the indentation depth of 800 nm for cells and 2-3 µm for spheroids.

### Rigidity index

To quantify the obtained differences between non-malignant and cancer cells in a manner independent of the culture and measurements conditions, we introduce the term rigidity index *R*:

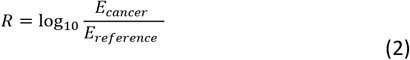

Rigidity index below < 0 indicates cell softening, while its value above zero denotes cell stiffening.

### Microrheological experiments in the frequency domain

The microrheological module applies a sine modulation to the piezoelectric scanner in the Z direction while being in contact with the cell (Fig. 1B, 1D). Broad oscillation frequencies can be applied during the measurements. They were 1, 5, 10, 25, 50, 75, and 100 Hz. Each frequency was acquired with 10 periods/frequency and a pause of 500 ms between frequencies. The amplitude was set to 50 nm, a constant sampling number of 600 obtained by adjusting the sample rate, Z-length of 6 µm, a set point of 2 nN and an approach velocity of 3 µm/s. The indentation depth varied between 1 – 2.5 µm. On average, 50 cells were measured for each cell line (between 40-60, depending on the cell line). Measurements were conducted in liquid conditions (i.e., cells in the corresponding culture medium) at RT and lasted no longer than two hours. Microrheological measurements were carried out above the cell nucleus, within the scan area of 9 µm^2^ (a grid of 3 pixels per 3 pixels was set). The measurements were repeated three times. The same parameters as above were applied for spheroids, except for the force limit (set to 15 nN), Z-length (15 µm). Here, the oscillation frequency varied from 1 Hz to 250 Hz. For each cell line, 15 spheroids were measured. On each spheroid, 2 maps were collected in the central part. The experiments were repeated two times.

### Determination of shear storage and loss moduli

Rheological properties of cells measured by AFM relied on the theoretical approach developed by Alcaraz et al. in 2003^37^. For small oscillation amplitudes, the complex shear modulus *E*\* for pyramidal indenter with a half open-angle *ϴ* is calculated can be described by the following equation:

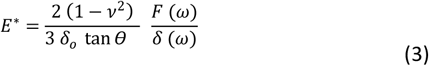

Where F is the load force and *δ* is the indentation depth. After applying Fourier transform and knowing the relationship between the elastic and shear moduli, i.e., *G = E/2(1+ν)*, the shear storage *G’* and loss *G”* moduli are:

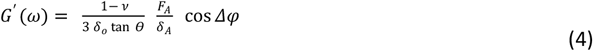

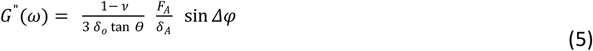

*G*^*′*^ *(ω)* measures the elastic energy that is stored and recovered during the oscillation, while *G”(ω)* stands for the loss modulus that considers the energy dissipated^49^.

### Power-law relations

The relations between *G*^*′*^ and *G”* plotted as a function of oscillation frequency were modelled with the power-law approach^32^:

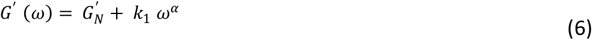

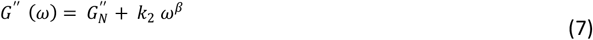

where *β* is the function of *α*. We used OriginPro to fit the data and Mathcad 2001 to determine the transition frequency, i.e. the crossover of *G*^*′*^(ω) and *G”(ω)*.

### Loss factor

The ratio between *G”* and *G*^*′*^ is called the loss factor. If its value is close to zero, a solid-like behaviour of the sample is assumed as there is no viscous contribution. A Newtonian fluid behaviour is assumed if the loss factor is close to 1 or above. The accuracy of the loss factor determination was quantified as a maximum error.

### Statistics

The following scheme was applied to nanomechanical and microrheological data to work on fully independent data sets. First, the means value was calculated for each elasticity map, and such data were used to create all plots. Thus, every single dot denotes a single elasticity map. In the case of cells, one elasticity map was recorded for one cell. Finally, the data are presented as a mean ± standard error of the mean (SEM). Statistical significance was obtained using a non-parametric Mann-Whitney test at the significance level of 0.05.

## Results

### Bladder cancer cells display cell- and culture-type dependent organization of actin filaments

Bladder cancer cell lines chosen for the study are: *(i)* non-malignant HCV29 cells derived from a non-malignant part of the irradiated, cancer-free region of the transitional bladder epithelium of the patient suffering from bladder carcinoma^50,51^; *(ii)* transitional cell carcinoma T24 cells derived from a female patient, aged 82, suffering from urinary bladder papillomatosis treated by electrocoagulation verified as urinary bladder carcinoma grade III^52^, and *(iii)* carcinoma HT1376 cells obtained from transurethral resection of invasive (grade 3) transitional cell carcinoma of 58-year-old Caucasian women^53^. As the first step of our analysis, the fluorescent images were collected for individual, separated cells and monolayers (**Fig. 2A,B**). The cell nucleus was labelled with Hoechst 33342, while the actin filaments were visualized using phalloidin conjugated with fluorescent Alexa Fluor 488 dye. Phalloidin binds specifically to the polymerized form of actin, i.e., F-actin, at the binding site located between F-actin subunits^54^. Therefore, this dye is widely applied to visualize filamentous forms of actin.

**Fig. 2:**
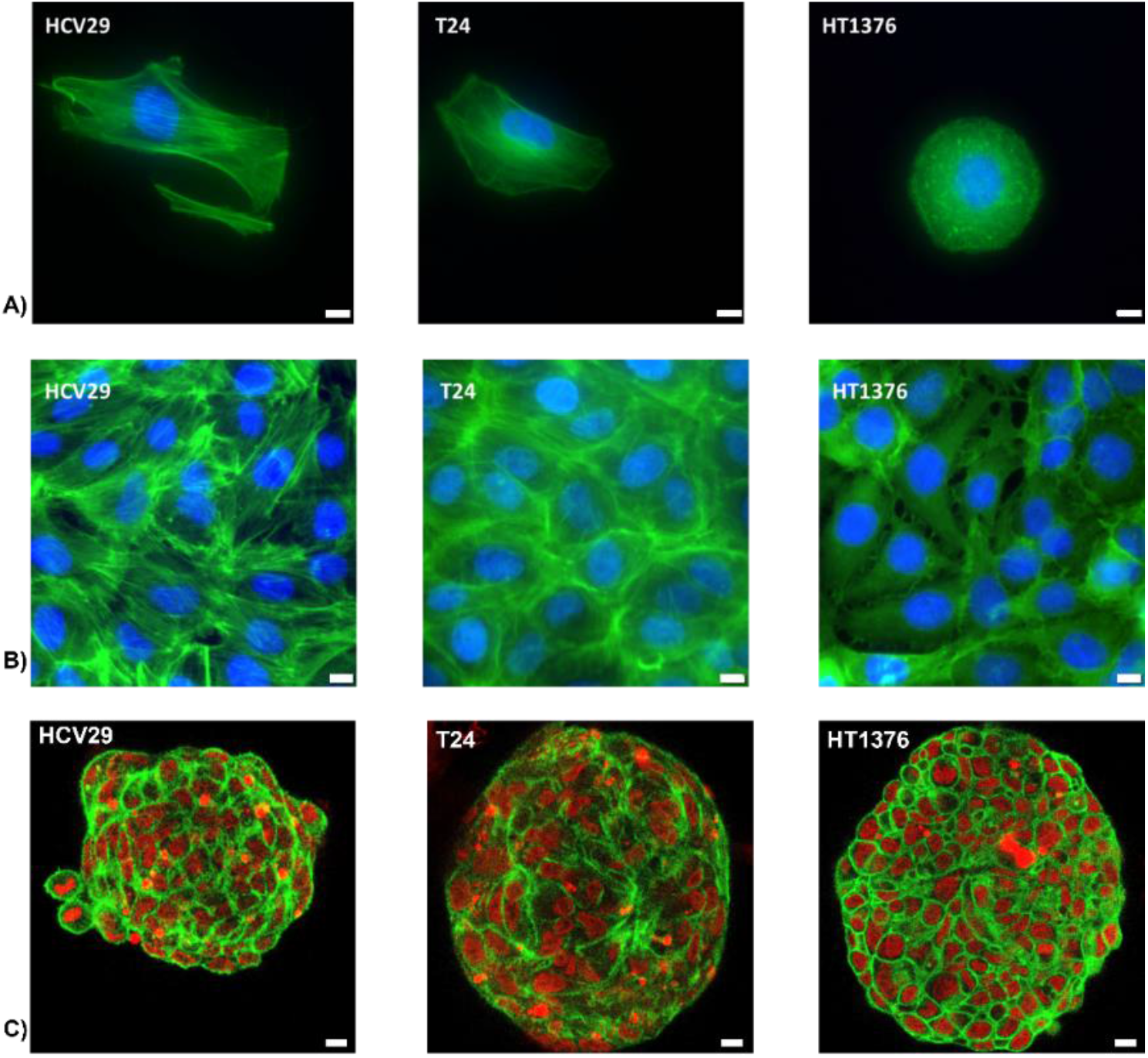
Fluorescent images showing the actin filaments organization inside bladder cancer cell lines at a different level of cellular organization. (A) single cells, (B) cell monolayers, (C) spheroids (one slice fron=m the Z-stack). For single cells, cell monolayers and spheroids, F-actin was stained using phalloidin – Alexa Fluor 488 (green), and In the case of single cells and monolayers, cell nuclei were stained with Hoechst 33342 dyes (blue)., while in the case of spheroids, cell nuclei were stained with SiR-DNA. Images were collected with the epifluorescence microscope (scale bar 10 µm), while spheroids were visualized using a confocal microscope (scale bar 10 µm).

Using an epi-fluorescent microscope, two ways of actin organization can be observed, i.e., long and relatively thick actin bundles that span over a whole cell or short F-actin filaments that form a shadow. HCV29 cells resemble the morphology of normal epithelium. They were elongated with clearly visible long and thick actin bundles spanning a whole cell. They are probably stress fibres. The number of actin bundles increases significantly when these cells form a monolayer. Cancer cells were different. T24 cells were more roundish with fewer long actin fibres than HCV29 cells. HT1376 cells did not show any actin fibres regardless of the culture conditions (individual cells or monolayer); thus, these cells appear to have a more homogeneous cytoskeletal structure. The thick actin bundles were more pronounced in HCV29 than in T24 cells and not detectable in HT1376 cells. This relation remains unaltered in both single cells and cell monolayers. Next, we ask how the actin cytoskeleton is organized in spheroids composed of the three studied cell lines. The actin cytoskeleton organization in spheroids is shown in the confocal images in **Fig. 2**. Actin filaments form a sheath surrounding the cell interior (mostly cell nucleus), inside which thick actin bundles were not visible (see also **Fig. 6**). The clarity of the actin sheaths is dependent on cell types. They are visibly in carcinoma HT1376 and non-malignant HCV29 cells. Sheaths are barely visible in transitional carcinoma T24 cells, but regions with brighter fluorescent signals denote high F-actin concentrations.

### G-actin monomers are predominant in cells lacking thick actin bundles

In cells, actin is present in monomeric G-actin and polymerized F-actin forms^55^. We hypothesize that the large amount of G-actin contributes to the viscous part of the cells. G- and F-actin expression was estimated from the Western blot by applying the densitometry approach (**Fig. 3**).

**Fig. 3.**
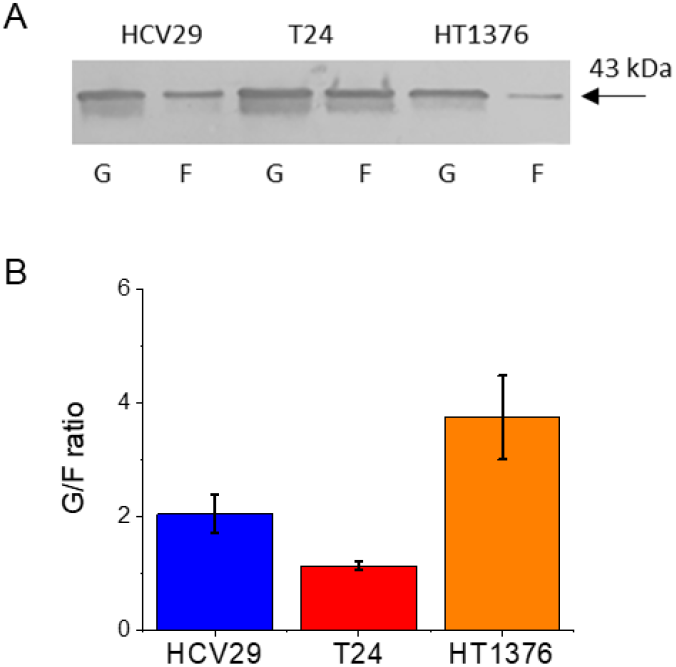
G/F actin ratio in bladder cancer cells. (A) Exemplary results from Western blot show total G- and F-actin expression in human bladder cells. The used antibody (against β-actin) targets all known actin isoforms with a molecular mass of 43 kDa (globular actin and fibrous actin). (B) Comparison of G/F ratio being smaller for cells displaying thick actin bundles (data are expressed as a mean ± SEM, from 3 to 4 independent repetitions).

A high G/F ratio denotes a large number of G-actin monomers compared to F-actin. Among the studied cell lines, the highest G/F ratio (3.74 ± 0.74) was obtained for HT1376 cells. It was 2-3 times higher than in the case of HCV29 and T24 cells, which indicated a greater level of G-actin monomers inside these cells. This correlates with the actin cytoskeleton organization characterized by actin mesh without thick actin bundles.

### Cancer-related softening of cells is independent of the cellular organization

Mechanical properties of individual bladder cancer cells followed the already published relation^27^; namely, the non-malignant HCV29 cells were more rigid than cancer (T24 and HT1376) cells. In this study, knowing that cells *in vivo* grow next to each other, we measured the mechanical properties of cells forming 2D monolayers. Fluorescent images, presented in **Fig. 2**, display various actin filaments organization with the highest, medium, and non-detectable amount of actin bundles in HCV29, T24, and HT1376 cells. AFM-based elasticity measurements showed that HCV29 cells are more rigid than cancer cells, regardless of the 2D culture conditions (**Fig. 4A**).

**Fig. 4:**
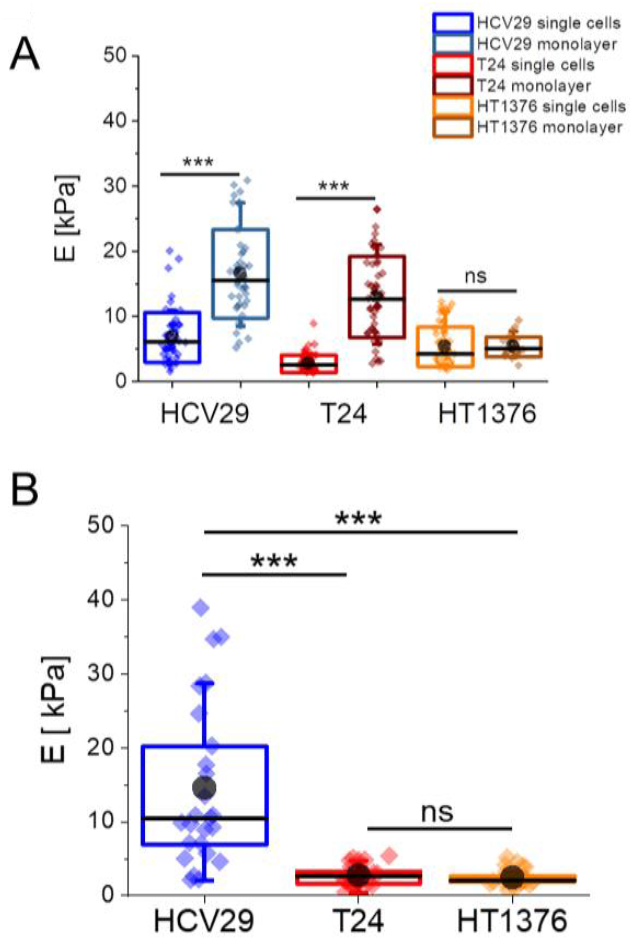
Differences in mechanical properties of bladder cancer cells in relation to culture conditions. (A) Box plots showing Young’s modulus variability for 2D cultured conditions, i.e., cell line, single cells or cell monolayer with MLCT-D cantilevers. Each point denotes the mean value calculated for single cells. The total number of cells measured was between 27 to 50 cells, depending on the cell type and culture conditions. The black dots and lines are the mean and median, respectively; box denotes standard deviation; error bars represent 10% and 90% percentile. (B) Analogous comparison between spheroids composed of normal and cancer cells, measured with NSC36-C cantilevers. For each cell line, 15 spheroids were measured (2 force maps/spheroid, each point denotes an individual elasticity map). (*** p < 0.001, ns – not statistically significant).

The apparent Young’s modulus of HCV29 (non-malignant cancer of ureter) cells cultured as a cell monolayer is larger than that of single cells. It increased from 6.8 ± 0.5 kPa (mean ± SEM; n = 55 cells measured) to 16.5 ± 1.0 kPa (n = 45 cells). A similar increment was observed for T24 (transitional cell carcinoma) cells. For these cells, elastic moduli raised from 2.7 ± 0.2 kPa (n = 64 cells) to 13.0 ± 0.9 kPa (n = 52 cells) for single cells and cell monolayer, correspondingly. On the contrary, the elastic modulus for HT1376 cells remained at the same level 5.3 ± 0.4 kPa (n = 58 cells) and 5.3 ± 0.3 kPa (n = 27 cells), regardless of the culture conditions. Obtained results indicate that the organization of actin filaments is strongly linked with the mechanical properties of cells. The role of stress fibres becomes larger in cells cultured as monolayers, showing a considerable modulus enlargement in cells with the well-developed actin cytoskeleton. Lack of stress fibres in carcinoma cells comes alongside unaltered mechanical properties.

Hereafter, we measured the mechanical properties of multicellular spheroids formed by the same bladder cell lines (**Fig. 4B**). It has been shown that the different seeding and growth conditions of the spheroids can affect their structure and physical properties^40^. Therefore, we used spheroids of 400 um diameter and after three days of culture in our measurements. In this way, we analyzed the spheroids from HCV29, T24 and HT1376 with a comparable size and originating from a similar culture day. As we would like to obtain the value that reveals the mechanics of cells embedded within the spheroid microenvironment, in the AFM measurements, NSC36-C cantilevers were applied as the tip is 13 microns long. This resulted in the indentation depth of 2-3 µm. Interestingly, we can observe that spheroids consisting of HCV29 cells were the stiffest with an elastic modulus of 14.5 ± 2.2 kPa (n = 25 maps recorded for 15 spheroids). Spheroids made of cancer cells were characterized by a modulus of 2.7 ± 0.3 kPa (n = 27 maps for 15 spheroids) and 2.4 ± 0.2 kPa (n = 28 maps for 15 spheroids), respectively for T24 and HT1376 cells. Notably, the mechanical properties of spheroids composed of cancer cell lines displayed similar mechanical properties. Altogether, these nanomechanical measurements show that cancer-related changes lead to softening regardless of the cells’ organization.

To compare measurements recorded with different types of cantilevers, we introduce the rigidity index *R* (defined as a moduli ratio of cancer to reference sample). The obtained *R* values confirm the softening of cancerous samples (**Fig. 5**).

**Fig. 5.**
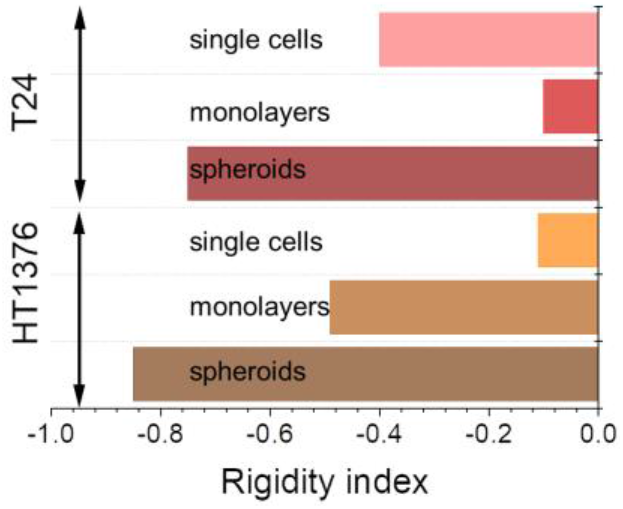
Rigidity index showing softening of cancerous samples. (single cell, monolayers and spheroids). The largest softening was observed for spheroids.

In the case of 2D cultures, cancer samples were softer than reference one; however, the degree of softening depended on the cell type. Nevertheless, the maximum softening was observed for the spheroids containing either T24 or HT1376 cells (R = -0.75 and -0.85, for T24 and HT1376, respectively).

### Fluid-like behaviour of cancer cells is observed at the single-cell and spheroid levels

Contact mechanics used to determine Young’s modulus assumes that living cells are purely elastic materials with high structural homogeneity. Cells are not fulfilling this assumption because they reveal viscoelastic nature^31,56^. Moreover, cell softening indicated by negative values of the rigidity index can be, supposedly, related to the larger fluidity of cancer cells^57,58^. Thus, in our next steps, we quantify the microrheological properties of the studied bladder cancer cells. Microrheological measurements were performed in the frequency domain (**Fig. 6**). The obtained storage *G’* and loss *G”* moduli were plotted as a function of oscillation frequency *ω*. Obtained relations, fitted with the power-law functions^32^, display the cell type and the culture conditions variability. A set of microrheology parameters was determined, i.e., storage *G*_*N*_’ and loss *G*_*N*_” shear moduli, loss factor, transition frequency *ω*_*T*_, and power-law exponent *α* **(Table 1)**.

**Fig. 6.**
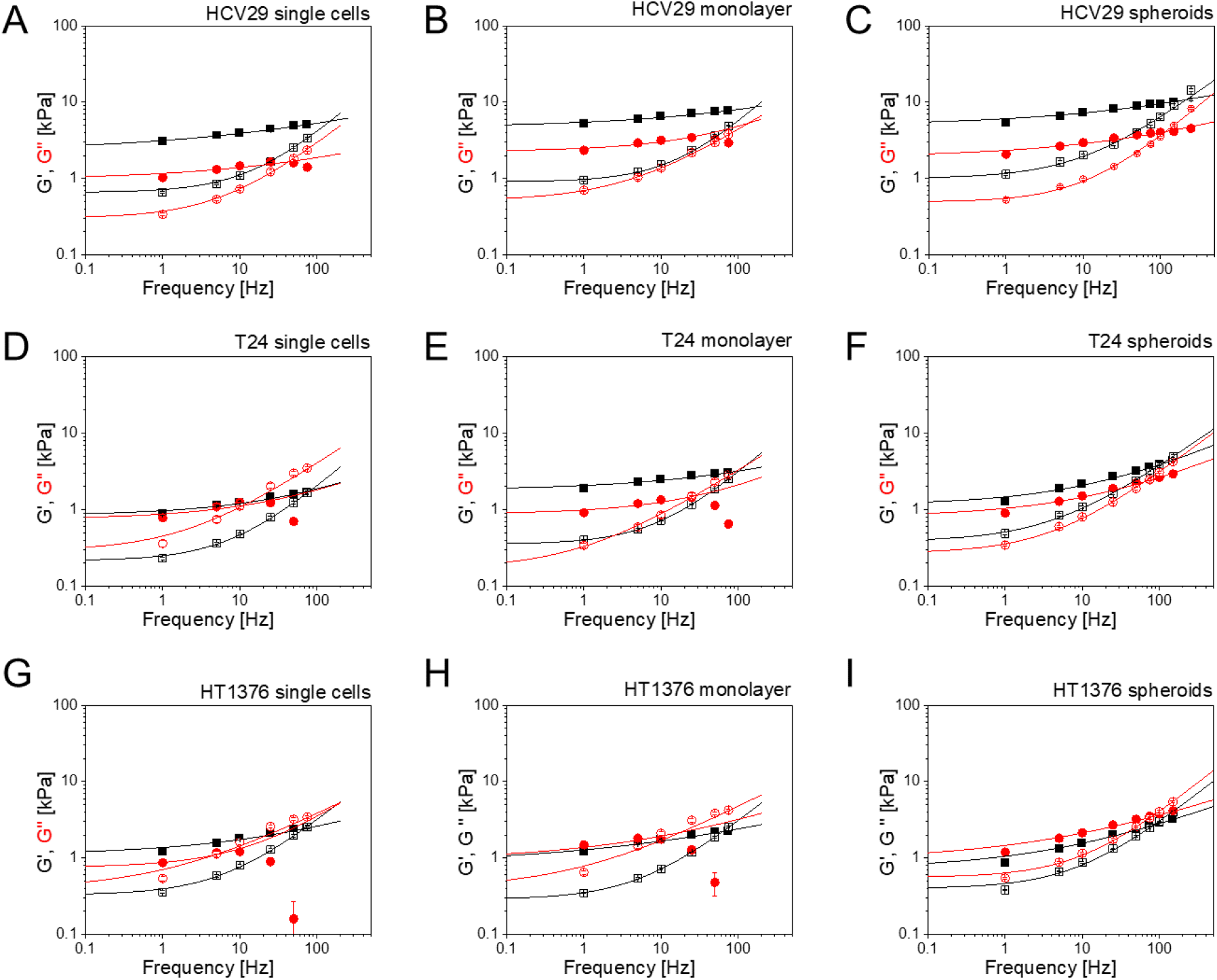
Microrheological properties of bladder cancer cells. Storage *G’* and loss modulus *G”* plotted as a function of oscillation frequency. The oscillation amplitude was 50 nm, and the frequency varied from 1-100 Hz for single cells- (A, D, G) and monolayers (B, E, H) and 1-250 Hz for the spheroids (C, F, I). Lines represent fits of the power-law function to the obtained data. Each point denotes the mean ± standard error of the mean from *n* = 30 to 40 cells and 15 spheroids from each cell line (2 maps/spheroid).

**Table 1.** Microrheological properties of bladder cancer cells. Storage *G*_*N*_’, loss *G*_*N*_” moduli and *α* were determined from the power-law fit. The error stems from the fit of the power-law function to the data. Loss factor is the ratio between *G*_*N*_”/*G*_*N*_’, while *ω*_*T*_ is determined from the intersection of *G’(ω)* and *G”(ω)*. The maximal error accompanies these quantities.

| Single cells | HCV29 | HCV29 Cyto-D | T24 | T24 Cyto-D | HT1376 | HT1376 Cyto-D |
| --- | --- | --- | --- | --- | --- | --- |
| $G_N'$ [kPa] | $2.27 \pm 0.12$ | $0.97 \pm 0.15$ | $0.84 \pm 0.06$ | $0.75 \pm 0.11$ | $1.14 \pm 0.10$ | $0.85 \pm 0.10$ |
| $G_N''$ [kPa] | $0.59 \pm 0.04$ | $0.24 \pm 0.01$ | $0.22 \pm 0.01$ | $0.29 \pm 0.14$ | $0.26 \pm 0.01$ | $0.39 \pm 0.07$ |
| loss factor | **$0.26 \pm 0.12$** | **$0.25 \pm 0.15$** | **$0.30 \pm 0.13$** | **$0.48 \pm 0.14$** | **$0.23 \pm 0.12$** | **$0.46 \pm 0.40$** |
| $\omega_T$ [Hz] | $143.2 \pm 0.3$ | $69.6 \pm 0.6$ | $89.6 \pm 0.4$ | $7.4 \pm 0.5$ | $70.0 \pm 0.7$ | $4.5 \pm 3.4$ |
| $\alpha @ G'(\omega)$ | $0.26 \pm 0.02$ | $0.58 \pm 0.21$ | $0.46 \pm 0.10$ | $0.46 \pm 0.30$ | $0.43 \pm 0.10$ | $0.71 \pm 0.06$ |
| Cell monolayer | HCV29 | HCV29 Cyto-D | T24 | T24 Cyto-D | HT1376 | HT1376 Cyto-D |
| $G_N'$ [kPa] | $5.03 \pm 0.26$ | $2.30 \pm 0.16$ | $1.79 \pm 0.13$ | $0.88 \pm 0.09$ | $1.14 \pm 0.13$ | $1.46 \pm 0.14$ |
| $G_N''$ [kPa] | $0.88 \pm 0.01$ | $0.51 \pm 0.04$ | $0.35 \pm 0.01$ | $0.17 \pm 0.04$ | $0.29 \pm 0.07$ | $0.49 \pm 0.14$ |
| loss factor | **$0.17 \pm 0.10$** | **$0.22 \pm 0.10$** | **$0.20 \pm 0.02$** | **$0.19 \pm 0.21$** | **$0.54 \pm 0.10$** | **$0.34 \pm 0.09$** |
| $\omega_T$ [Hz] | $165.8 \pm 1.6$ | $141.5 \pm 2.3$ | $100.7 \pm 0.1$ | $26.6 \pm 1.3$ | $60.6 \pm 0.7$ | $13.6 \pm 3.5$ |
| $\alpha @ G'(\omega)$ | $0.41 \pm 0.10$ | $0.54 \pm 0.23$ | $0.36 \pm 0.09$ | $0.52 \pm 0.25$ | $0.35 \pm 0.10$ | $0.58 \pm 0.75$ |
| Spheroids | HCV29 | HCV29 Cyto-D | T24 | T24 Cyto-D | HT1376 | HT1376 Cyto-D |
| $G_N'$ [kPa] | $4.99 \pm 0.38$ | $1.92 \pm 0.16$ | $1.16 \pm 0.09$ | $0.81 \pm 0.07$ | $0.77 \pm 0.08$ | $1.06 \pm 0.12$ |
| $G_N''$ [kPa] | $1.01 \pm 0.07$ | $0.49 \pm 0.04$ | $0.38 \pm 0.04$ | $0.27 \pm 0.03$ | $0.29 \pm 0.03$ | $0.44 \pm 0.03$ |
| loss factor | **$0.20 \pm 0.15$** | **$0.26 \pm 0.10$** | **$0.33 \pm 0.07$** | **$0.33 \pm 0.20$** | **$0.38 \pm 0.25$** | **$0.42 \pm 0.06$** |
| $\omega_T$ [Hz] | $214.8 \pm 1.9$ | $116.4 \pm 1.0$ | $106.7 \pm 0.9$ | $62.8 \pm 0.7$ | $79.5 \pm 0.7$ | $73.9 \pm 0.9$ |
| $\alpha @ G'(\omega)$ | $0.31 \pm 0.04$ | $0.34 \pm 0.06$ | $0.47 \pm 0.05$ | $0.46 \pm 0.05$ | $0.46 \pm 0.05$ | $0.44 \pm 0.06$ |

Storage modulus *G*_*N*_′, related to the elastic energy stored in the measured sample, describes a storage shear modulus *G*^*′*^ value at *ω* = 0. It is the highest for single non-malignant HCV29 cells, indicating higher shear forces resistance. Lower storage modulus obtained for cancer cells indicated larger cellular deformability induced by shear forces. It correlates with Young’s modulus (compression of cells). Analogously, the loss shear modulus *G*_*N*_”, which corresponds to the dissipated energy in the sample, describes *G”* value at *ω* = 0, and it is lower than *G’* for low frequencies. It is larger for non-malignant and lower for cancerous ones, and it seems to be independent of the cellular organization. The dissipative nature of loss shear modulus shows that the viscous component cannot be neglected.

Cellular viscoelasticity results from the water content and polymerized structural elements. This means that cells behave either like solid-like or fluid-like materials, both in the frequency and timescale domains. Thus, the loss factor was calculated to assess the behaviour of non-malignant and cancerous cells. The loss factor equals 0.21 ± 0.05 (n = 3, i.e., loss factors for single cells, monolayers and spheroids), 0.28 ± 0.07 and 0.38 ± 0.16, in order for HCV29, T24 and HT1376 cells when averaging over data, regardless of the culture conditions (2D cultures or spheroids). A higher loss factor suggests a shift towards the fluid-like behaviour of the cells. The detailed analysis of the obtained loss factors indicates that the presence and organization of thick actin bundles inside the cells barely correlate with the loss factor at the level of single cells. It is equal to 0.26 ± 0.12, 0.30 ± 0.13, 0.23 ± 0.12 for non-malignant HCV29 and cancerous T24 and HT1376 bladder cells, respectively. Considering the maximal error calculated here, we can state that the dissipative component remained similar for all studied cell lines at the single-cell level. Contacts with neighbouring cells (monolayers) or their embedment in ECM (spheroids) contribute to the loss factor changes in a cell-dependent manner. Its value is the largest for carcinoma HT1376 cells and the smallest for non-malignant HCV29 cells. The loss factor is correlated with the power-law exponent, which measures the proximity of the experimental data to the fluid-like or solid-like behaviour of the studied material. One would expect that the loss factor calculated from the power-law exponent, as *tan((π/2)·α)*, would depend linearly on the loss factor determined as a ratio *G”/G’*. Even so, in our case, such behaviour was not observed, which probably stems from two main reasons. First, *G’ (ω) c*annot be fully modelled as a single power law. Second, *G*_*N*_’ and *G*_*N*_” value are obtained from the fitted power-law model at *ω* = 0. When considering the power-law exponent, obtained in such a way, the fluid-like behaviour in the studied cells is observed at the single-cell and spheroid levels.

### Cytochalasin D reveals the link between the presence of actin fibres and the storage modulus, but not the loss modulus

To elaborate the role of actin filaments in maintaining the rheological properties of bladder cancer cells, the cells from 2D cultures and spheroids were treated with 5 µM cytochalasin D (cyto D). Cytochalasin D is a permeable toxin that impairs actin filament organization by inhibiting the polymerization of actin filaments^59^. The effect of cytochalasin D on mechanical properties is also known for bladder cancer cells^27,60^; therefore, we focused on evaluating its effect on the rheological properties of cells. The first observation is that cells in 2D cultures can be more efficiently detached from the surface by applying shear forces (**Fig 6**, red dots). Regardless of the cell type (HCV29, T24, and HT1376), a drop of storage modulus appears above 10-20 Hz. As the actin cytoskeleton is disrupted, individual cells lose their attachment to the surface, leading to lower resistance to shear forces and probably easier detachment; despite this, the full detachment of cyto D treated cells was not visible under AFM. Such effect is not observed in spheroids because it is more challenging to remove single cells from a 3D environment even for high frequencies (maximum applied modulation frequency was set to 250 Hz). Simultaneously, the effect of cyto D on spheroids was clearly visible in confocal images of spheroids formed of each bladder cancer cell line used in our study (**Fig. 7**). Loss factor and power-law exponent obtained from all the studied bladder cancer cells show a considerable variability that is difficult to be correlated with changes in actin cytoskeleton before and after treatment with cyto D (**Table 1**). Looking at the storage modulus of cyto D treated cells, a drop by 20% to 40% only for HCV29 and T24 cells was detected. In these cells, thick actin bundles are seen. The storage modulus of HT1376 cells decreased only for single cells, but it increased for cyto D treated cells that grow as a monolayer and in spheroids. Such behaviour seems to be linked with a lack of thick actin bundles in these cells. Loss modulus dropped from 0.59 ± 0.04 kPa to 0.24 ± 0.01 kPa, from 0.88 ± 0.01 kPa to 0.51 ± 0.04 kPa, and from 1.01 ± 0.07 kPa to 0.49 ± 0.04 kPa for non-malignant HCV29 cells, regardless of the culture conditions. The drop of loss modulus is a sign of the low dissipative nature of the studied cells. On the contrary, HT1376 cancer cells present an increase in loss modulus, denoting the stronger dissipative characteristic of these cells.

**Fig. 7.**
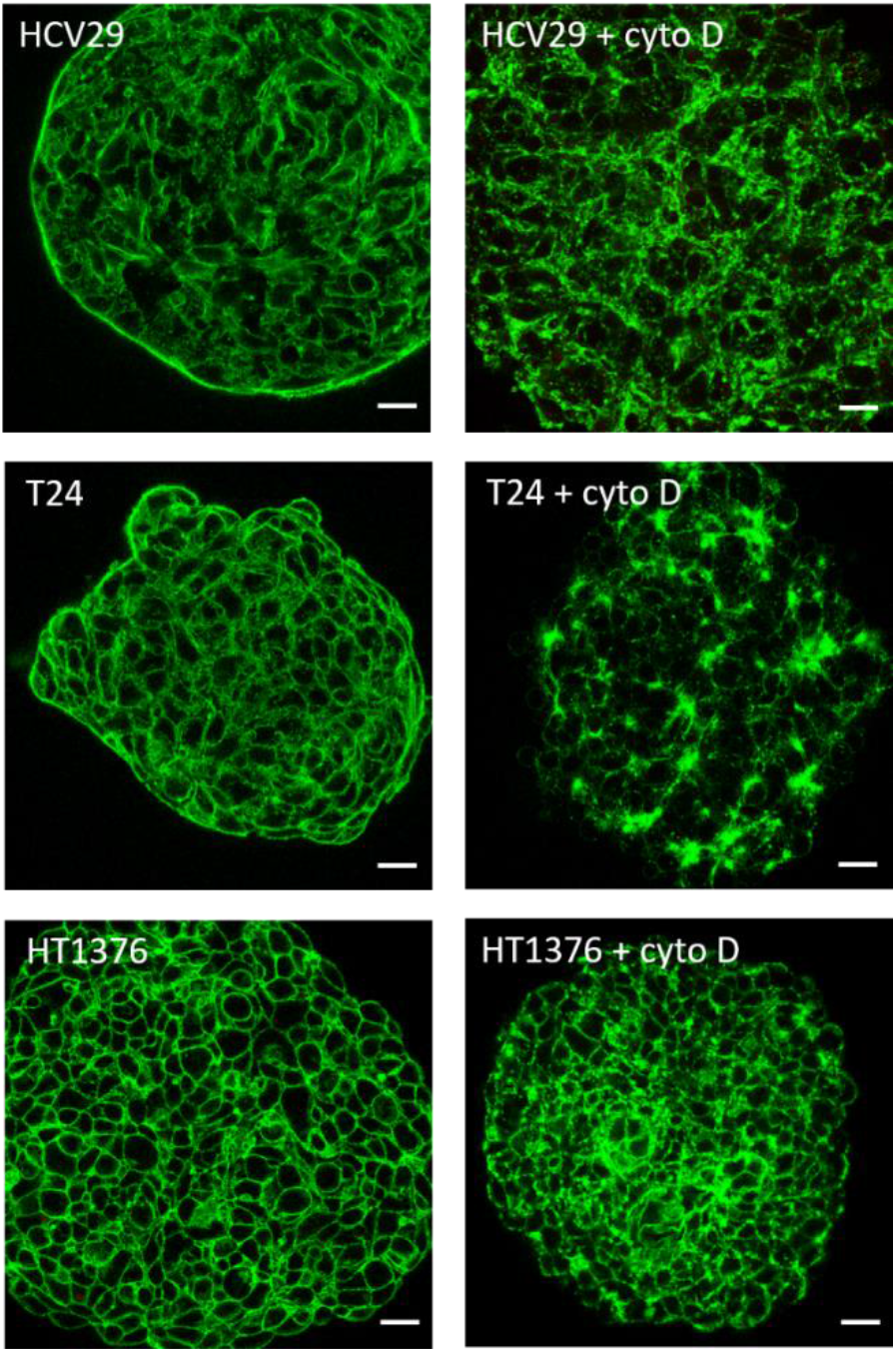
Cytochalasin D prevents polymerization of F-actin in spheroids. Confocal image of F-actin stained with phalloidin conjugated with Alexa Fluor 488 dye. Scale bar = 10 µm

These results can be associated with the cytoskeleton organization inside these cells, well-differentiated in non-malignant HCV29 cells and poorly differentiated in carcinoma HT1376 cells. When looking into the transitional cell carcinoma T24 cells, an increase of the loss modulus for single cells indicates that they are close to HT1376 cells, while the decrease for cell monolayers and spheroids may result from the extensive formation of kind of sheaths built up from actin filaments. Therefore, we can affirm that the loss factor correlates with the actin cytoskeleton structure of the cells.

### Fluid-like behaviour of cancer cells is identified with a lower transition frequency and a higher power-law exponent

Next, we asked whether we could detect the transition between solid-like and fluid-like behaviour of cells treated with cyto D. Thus, we analysed the transition frequency *ω*_*T*_ and compared it to the power-law exponent (**Fig. 8, Table 1**). The obtained results show that the transition frequency decreases for cancer cells, regardless

**Fig. 8.**
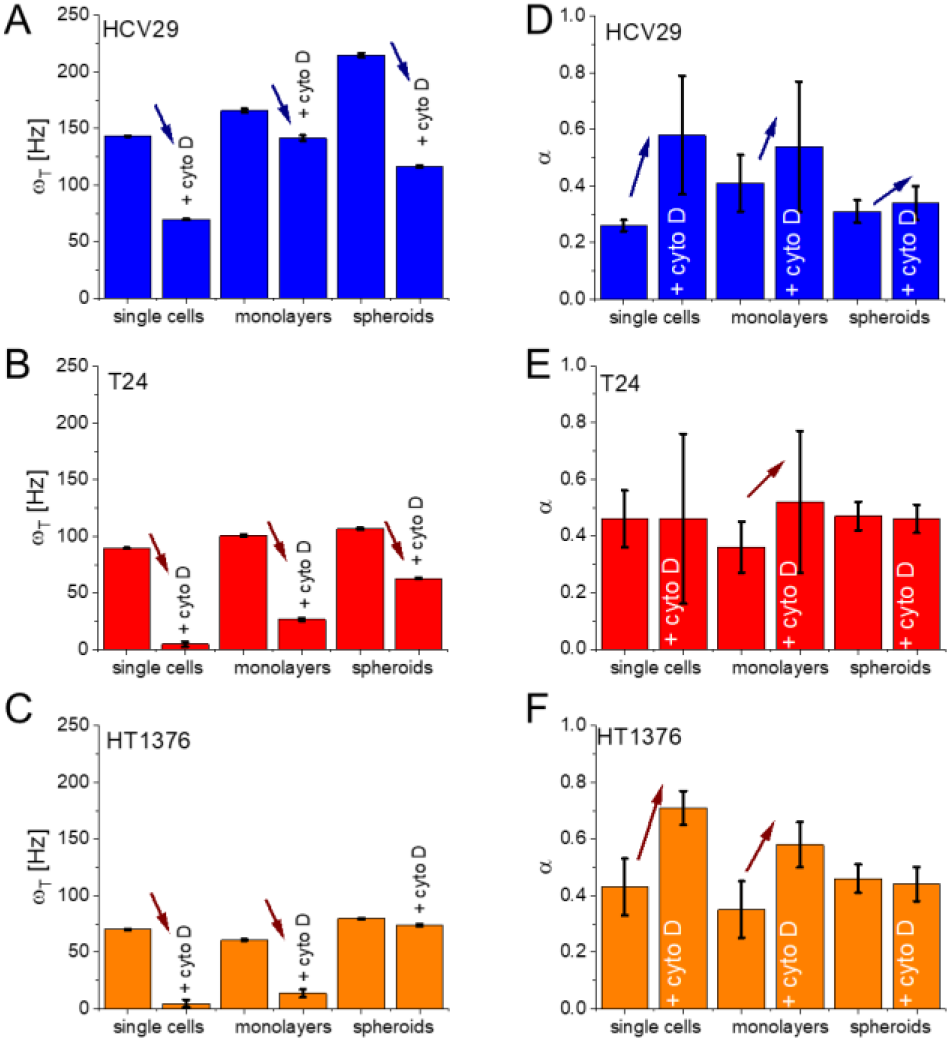
Transition frequency and power-law exponent changes with the altered actin cytoskeleton. Transition frequency for non-malignant (A) and cancer cells (B,C) treated with cyto D plotted as a function of culture conditions. (D-F) The analogous plot of power-law exponent derived from the fit to *G’ (ω)* data.

**ARTICLE Journal Name** of the culture conditions. For single cells, it dropped from 143.2 ± 0.3 Hz (for non-malignant HCV29 cells) to 89.6 ± 0.4 Hz and 70.0 ± 0.7 Hz for T24 and HT1376 cells, respectively. The corresponding relations for the *ω*_*T*_ decrease were *(i)* 165.8 ± 1.6 Hz (HCV29 cells) to 100.7 ± 0.1 Hz (T24 cells) and 60.6 ± 0.7 Hz (HT1376 cells) and (*ii)* 214.8 ± 1.9 Hz (HCV29 cells) to 106.7 ± 0.9 Hz (T24 cells) and 79.5 ± 0.7 Hz (HT1376 cells) for cells cultured as monolayers and spheroids. Thus, the lower transition frequency is characteristic for cancer cells^31,61^. Power-law exponent for non-malignant HCV29 cells equals to 0.26 ± 0.02 (single cells), 0.41 ± 0.10 (monolayers), and 0.31 ± 0.04 (spheroids). The analogous relations for cancer cells were 0.46 ± 0.10, 0.52 ± 0.25, 0.46 ± 0.05 for transitional cell carcinoma T24 cells and 0.43 ± 0.06, 0.35 ± 0.10, 0.46 ± 0.05 for HT1376 cells. The means of the power-law exponent for each cell lines are 0.35 ± 0.05 (n = 3 culture conditions), 0.43 ± 0.06 (n = 3), 0.41 ± 0.06 (n = 3) for HCV29, T24 and HT1376 cells, respectively. Its larger value indicates a shift towards fluid-like behaviour of cancer cells. This matches with the low transition frequency obtained for cancer cells.

To evaluate the contribution of actin filaments in the fluidization of cells, transition frequency and power-law exponent were determined for cyto D treated cells. The transition frequency for cyto D treated cells becomes significantly lower than the untreated cells, except in the case of HT1376 cells that form spheroids. Moreover, the treatment with cyto D alters the power-law exponent, too. For non-malignant HCV29 cells, this value raised at the single-cell and cell monolayer level, showing an increase in fluid-like behaviour of these cells only in 2D cultures (**Fig. 8**). In the case of cancerous T24 cells, the power-law exponent in cyto D treated cells increase only in cells cultured as monolayers where this toxin damaged the enhanced number of thick actin bundles. In the case of the cancerous HT1376 cells, the cyto D treatment increased power-law exponent only in 2D cultures but not in spheroids.

Altogether, we can conclude that the fluid-like behaviour of cancer cells is identified with a lower transition frequency and a higher power-law exponent.

## Discussion

A living cell has a dynamic structure that changes in response to its functioning and various microenvironmental conditions. The viscoelastic nature of living cells stems from their internal structure, in which both fluid and solid phases can be distinguished. The viscoelasticity of cells results from the water content (i.e. cell cytosol contains most of the fluid cellular constituents) interlaced with the polymerized solid structures (such as cell cytoskeleton). Therefore, as previously reported in numerous papers, the mechanical and rheological properties are extremely complex and challenging to be fully understood^62,63^.

Our work is mainly focused on the nanomechanical and rheological properties of bladder cancer cells. The measurements of Young’s modulus by AFM have already been used as a relevant method to distinguish the cancerous cells by exploring the process of their cytoskeletal reorganization^22,64^. It has been demonstrated that the mechanical properties of cells are mainly dependent on the cytoskeleton structure, particularly on actin filaments, which are known to behave like viscoelastic materials^65,66^. Thus, both elastic and viscous components must be considered to obtain a complete description of cell mechanics^67^. In our studies, we chose three bladder cancer cell lines that display different actin cytoskeleton organisations and diversely behave when grown as 2D cultures (single cells and monolayers) and 3D spheroids. We used the AFM to characterize their mechanical and rheological properties. In several works, the bladder cancer cells have already been measured in nanoindentation experiments, showing that the non-malignant cells are stiffer than the cancer cells^25–27,68^. However, those measurements were conducted only at the single-cell level, and the rheological properties of these cells were not extensively studied. In two published papers prior to this, the rheological properties of some bladder cancer cells (RT112, T24 and J82) were compared, suggesting that changes in cell rheology may be a signature of cancer metastasis and that it correlates with different invasiveness of bladder cancer^32,61^. The studied carcinoma RT112 bladder cancer cells are known to be moderately differentiated, whereas the transitional cell carcinoma T24 and J82 cells are poorly differentiated and have a higher malignancy potential.

In our experiments, to compare the nanomechanical and rheological properties of cells, we decided to use another panel of bladder cancer cell lines: one originating from early (non-malignant cell cancer of ureter, HCV29 cells), medium (bladder carcinoma, HT1376 cells) and late (transitional cell carcinoma, T24 cells) stages of cancer progression. These cells are characterized by a different organization of the actin cytoskeleton, which we assume to be a key player in their physical properties. The chosen cell types have already been measured by AFM, as single cells^25,27,45,60^. It has been demonstrated that the differences in their mechanical properties are primarily linked with the organization of actin filaments. The nanomechanical results exhibited the highest Young’s modulus for non-malignant HCV29 cells and the lowest for transitional cell cancer T24 cells. Young’s modulus of carcinoma HT1376 cells was ranked between them. Our previous research^27^ demonstrated that the content of β-actin in cancerous cell lines was 35% and 42% in respect to HCV29 cells, respectively, for HT1376 and T24 cells. Those results suggested that when single cells with increasing tumour grading are measured, the actin content decreases in the cancerous cells compared to non-malignant cells.

The present study detects the nanomechanical and rheological properties as a function of culture conditions, i.e., 2D cultures (single cells and monolayers) and spheroids. In parallel, we quantified the amount of unpolymerized G-actin, supposing that its larger expression contributes to the viscous component of the cell. The results show that cancer cells are softer than non-malignant cells indicating their large deformability. This is in good agreement with the previous results^27^. Although the correlation between cell mechanics and the expression of G-actin is not obvious, when considering the mechanical properties of cells in monolayers and spheroids, the largest deformability is observed for carcinoma HT1376 cells, for which the amount of G-actin is 2-3 times larger than in the other cells. Consequently, the cell and spheroid softening can result from low actin content and significant G-actin expression. In addition, the data indicate that a low abundance of actin filaments is characteristic for the mechanics of cancer cells. Notably, we found out that the culture conditions (2D cultures or spheroids) did not influence the mechanical properties of bladder cancer cells. Indeed, the non-malignant HCV29 cells remained stiffer than cancer cells regardless of the culture conditions making Young’s modulus a potential biomechanical marker of bladder cancers. Nevertheless, it should be underlined here that the distinction between HT1376 and T24 cells through nanomechanical properties is rather difficult because, at the single-cell and spheroid levels, both cancer cell lines reveal similar mechanical properties. It is known that monolayer cells exchange mechanical and molecular signals with the adjacent cells, which would be the reason behind the changes in their cytoskeleton organization^69^. We had observed that HCV29 and T24 cells increased their stiffness when cultured as monolayers, while HT1376 preserved mechanical properties in both conditions. For these reasons, we could associate the observed difference in cell mechanics with the presence and absence of thick actin bundles.

Nanomechanical properties are typically related to resistance to compression. Recently, it has been reported that cancerous cells during invasion become more fluid, which may help them for distant metastasis^70,71^. We applied the sinusoidal oscillations at specific indentations to probe the rheological response of the cells^37^. The results show that HCV29 and T24 cells stiffen with increased cellular organisation complexity. The storage modulus is bigger in the case of spheroids than that obtained for single cells. On the contrary, HT1376 cells become softer. Therefore, we can affirm that the shear storage modulus follows the same trend as Young’s modulus. This clearly indicates that thick actin bundles contribute to the mechanical resistance of cells undergoing shear stress. Moreover, we suggest that the storage modulus can be used interchangeably with Young’s modulus as a potential biomechanical marker of bladder cancers. Both Young’s and storage modulus describe only the elastic component of cells induced by deformation forces (here, compression and shear). Thus, we can declare that non-malignant HCV29 cells are more resistant to such deformations than cancerous T24 and HT1376 cells. Importantly, the evaluation of elastic changes through rigidity index showed that the largest cancer-related softening induced by compression occurred in the cancer spheroids, not in 2D cultures. The opposite effect was observed when shear forces induced the elastic deformation. In this case, the largest softening occurred for single cancer cells.

As we remarked before, microrheological measurements can give information about the dissipative component in the mechanical properties of living cells. It can be described using different parameters. The simplest is loss modulus, which for the studied bladder cancer cells constitute a significant percentage from 20% to 50% of the storage modulus, depending on the cell type and culture conditions. This means that the viscoelastic component in the bladder cancer cells plays a key role in defining their mechanical properties, and it cannot be ignored. Moreover, cells are dynamic systems; thus, they change their mechanical properties in response to various stimuli, including external mechanical forces. More precisely, compressive and shear stress can drive cells towards a more invasive phenotype depending on their magnitude^62,63^. They may induce a transition from a solid-like to fluid-like behaviour. The latter one has been attributed to the invasiveness of the cancer cells^64,65^. Likewise, by applying oscillatory modulations with increasing frequency, the mechanically induced fluidization of cells can be studied by analysing the power-law exponent and the transition frequency. Our results show that HCV29 has the highest value of the transition frequency, compared to T24 and HT1376 cells, regardless of the culture conditions. Additionally, in these non-malignant HCV29 cells, the transition frequency increases with the organization level from single cells to spheroids. A similar but smaller increase was observed for T24 cells as well. We could explain these effects with the presence of thick actin bundles in these cells. Unlike HCV29 and T24, the carcinoma HT1376 cells have a different cytoskeletal structure, and consequently, the absence of stress fibres does not modify the viscoelastic properties when the confluency is altered. A significant expression of monomeric G-actin could support this effect. Finally, to reinforce these results, experiments with cytochalasin D were performed. Upon inhibiting the polymerization of actin filaments with cyto D, we observed that in 2D cultures, the transition frequency decreased in all cells and that they showed a more fluid-like behaviour. A similar trend was observed in other studies in which they used different compounds to disrupt the actin cytoskeleton ^32,66^. Interestingly, in our current study, we show that the oscillatory measurements could be performed at higher frequencies when we use the multicellular spheroids, which we suggest as a better model than the 2D cultures. From the confocal images, we could see differences in the actin cytoskeletal organization between treated and untreated spheroids. And what is relevant for us is that, unlike HCV29 and T24, HT1376 cells had an increase in loss modulus in cyto D treated spheroids, implying the stronger dissipative peculiarity of these cells again.

## Conclusions

Our study demonstrated that to obtain a complete picture of the viscoelastic properties of bladder cancer cells, both nanomechanical and rheological measurements must be done. We compared a panel of bladder cancer cell lines originating from early (non-malignant cell cancer of ureter, HCV29 cells), medium (bladder carcinoma, HT1376 cells) and late (transitional cell carcinoma, T24 cells) stages of cancer progression. Measurements were conducted for single cells, monolayers and spheroids. Our findings show that both Young’s and storage modulus are greater in non-malignant HCV29 cells, indicating their larger resistance to compression and shear forces than cancer cells. The evaluation of elastic changes through rigidity index showed that the substantial cancer-related softening occurred in the spheroids for compressive forces and in single cells for shear forces. Rheological measurements demonstrated that the viscoelastic character of cells could not be neglected. The loss modulus was the lowest for single cells and increased for monolayers and spheroids in a cell type-dependent manner. Treatment with cyto D revealed the role of actin filaments in maintaining the mechanical properties of cells possessing a well-organized cytoskeleton with the existence of thick actin bundles (here, HCV29 and T24 cells). For HT1376 cells, in which thick actin bundles were present neither in single cells nor in monolayers, actin filaments are not the dominant players in the mechanical properties of cells. Fluid-like behaviour was identified by analysing transitions frequency (indicate fluidity when decreases) and by the power-law exponent (indicate fluidity when increases). The power-law exponent was independent of the culture conditions (2D culture, spheroids); however, it showed that cancer cells shift towards more fluid-like behaviour. The transition frequency increased for HCV29 and T24 cells as a function of cellular organization, inducing solid-like behaviour of more complex cellular systems in cells displaying thick actin bundles in the actin cytoskeleton. For cells with the poorly differentiated actin cytoskeleton (HT1376 cells), transition frequency oscillated within 60 Hz – 79 Hz regardless of the cell organization level. The transition frequency varied in a specific manner for each cell line, making it a potential tool to classify the cancer type. In conclusion, we have confirmed that the viscoelastic properties of the cells could be used as a marker of malignancy and that the rheological characteristics can be associated with the rearrangement and organization of the cell cytoskeleton.

## Author Contributions

KG, ML designed and discussed the experiments, KG measured and analyzed the data, SKK participated in the G/F actin determination, while JP participated in Western blot and fluorescent microscopy. KG, ML wrote the manuscript and prepared all figures. All authors contributed to results interpretation and read the manuscript.

## Acknowledgements

This work is supported by the European Union under the Marie Skłodowska-Curie grant agreement No. 812772 (Phys2BioMed).

